# Fibril fragmentation generates diversity in seed population

**DOI:** 10.1101/2024.11.12.623277

**Authors:** Júlia Aragonès Pedrola, Marija Todorovska, Shalini Singh, Assaf Friedler, Stefan G. D. Rüdiger

## Abstract

Neurodegenerative diseases are characterised by the formation and accumulation of protein fibrils. The mechanism underlaying this aggregation process remains poorly understood. Fibril fragmentation, resulting in seed generation, plays a role in toxicity. Here we provide a quantitative picture of the impact of ultrasound on patient-derived and recombinant fibrils from various diseases. Fragmentation of recombinant Tau fibrils and patient-derived fibrils from Alzheimer’s Disease, Corticobasal Degeneration and Frontotemporal Dementia generates amyloid and non-amyloid species. Interestingly, patient-derived fibrils are more susceptible to ultrasound than artificial fibrils. Understanding fibril fragmentation and the generation and nature of seeds may provide insights to the molecular mechanism of the disease progression, contributing to the development therapeutic approaches.

## Introduction

Neurodegenerative diseases, such as Alzheimer’s Disease (AD), Parkinson’s Disease (PD) and Huntington’s Disease (HD), are characterized by progressive deterioration of neuronal structure and function, resulting in cognitive decline, motor impairment and neurological symptoms^1^. Currently, disease-modifying therapies targeting the casual mechanism are emerging^2, 3^. However, the mechanism underlying neurodegenerative diseases is poorly understood.

A common feature of these diseases is the accumulation of protein fibrils ^4–6^. Understanding the mechanisms of fibril formation and propagations are key for a comprehensive picture of the origin at molecular level. Recently, seeds resulting from fibril fragmentation are hypothesized to be involved in cell-to-cell spreading ^7–9^. Protein aggregation chemistry has contributed to getting insights into the aggregation process. Fibril formation starts with a conformational change of monomer, named primary nucleation. Upon the addition of monomers, the aggregation further grows through an elongation phase, resulting in fibril formation. Fibrils can undergo fragmentation, leading to the formation of seeds. In this secondary nucleation, the seeds, which have elongation-competent fibril ends, are able to accelerate the amyloid formation by-passing primary nucleation ^10^.

Fibril fragmentation depends on their stability, which determines the propensity of fibrils to break. An established technique to induce fibril fragmentation is application of ultrasound via sonication^11–16^. Unstable fibrils, prone to fragmentation, are more susceptible to the generation of seeds. In consequence, an enhanced seed formation, can ultimately contribute to toxicity and disease progression^7–9^. Characterization of fibril fragmentation and the products resulting from are crucial for the rational development of therapeutic strategies, by stabilizing fibrils or target seeds.

Here, we determined the half-breaking time of patient-derived Tau fibrils from AD, Corticobasal Degeneration (CBD) and Frontotemporal Dementia (FTD) and recombinant repeat domain fibrils exposed to ultrasound. We demonstrated that all fibrils are susceptible to sonication. Interestingly patient-derived fibrils, characterised by larger fibril diameter, fragment at a higher rate than the narrower artificial fibrils. Importantly, we identified that sonication generates a heterogeneous population, composed of amyloid and non-amyloid species.

## Results

### Seeds induce TauRD aggregation

Tau aggregation characterises the development of AD and other tauopathies, including CBD and FTD^17, 18^. Here we use the Tau Repeat Domain (Q244-E372, TauRD) with pro-aggregation mutation ΔK280. Approaches to induce Tau aggregation consist of the addition of heparin, which promotes the conformational change in the monomer, or seeds. To confirm the ability of seeds to induce aggregation we performed homologous seeding using TauRD monomers and seeds and compare it to heparin-induced aggregation. We used seeds generated by direct probe sonication of pre-formed TauRD fibrils, for either five or 30 seconds, 40% of amplitude and 750 Watt. We performed Thioflavin-T (ThT) aggregation assay to monitor fibril formation. ThT is a fluorescent dye which emits fluorescence upon binding to beta-sheet structure, increasing fluorescent emission during fibril formation^19^.

All conditions resulted in an increased ThT fluorescence emission (Fig A-C), indicating fibril formation. Tau aggregation induced by seeding resulted in an accelerated amyloid formation, as observed by an earlier saturation of the fluorescent signal. Interestingly, five- and thirty-seconds seeds show a similar aggregation curve, indicating that the nature of seeds does not affect the aggregation reaction.

**Figure 1.**
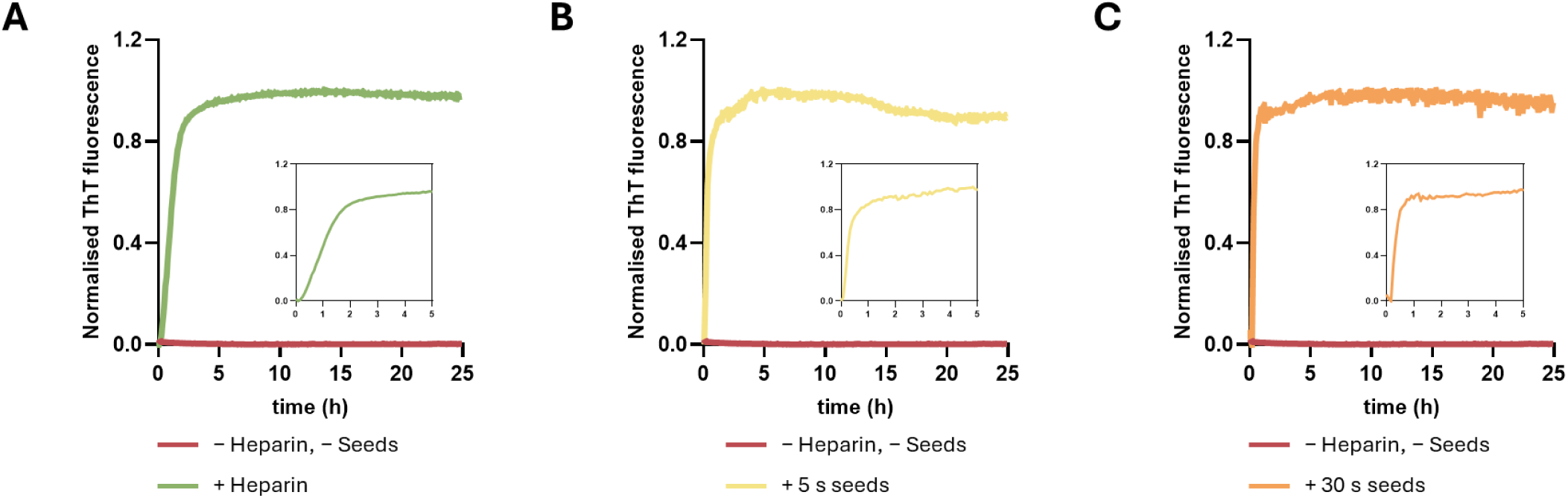
Tau aggregation is induced by seeds. ThT aggregation assay of 20 μM TauRD induced by (A) 5 μM heparin, (B) 5 μM seeds, 5 seconds sonication and (C) 5 μM seeds, 30 seconds sonication.

### Ultrasound reduces recombinant fibril size in a time-dependent manner

To quantify the impact of ultrasound on amyloid fibrils, we aimed to quantify sonication-dependent size changes throughout a time course. To do so, we used the Fibril Paint/FIDA assay (Fig. 2A). Fibril Paint are a class of peptides that specifically bind in a non-covalent manner to amyloid fibrils ^20, 21^. Fluorescently labelled Fibril Paints are thus suitable to prepare fibrils for quantitative, fluorescence bases assays, such as Flow Induced Dispersion Analysis (FIDA). The FIDA assay measures the size-dependent dispersion of fluorescently labelled molecules in solution ^22, 23^. As a read-out it provides the hydrodynamic radius, R_h_, a parameter corresponding to the mean size of biocomplexes in solution. Here we use FibrilPaint6 [`(Fl)-PWWRRPWWPWHHPH`], a 14 residue and fluorescein (Fl) to visualise the fibrils throughout this study ^21^. FibrilPaint6 bound to amyloid fibrils disperses slower in FIDA, resulting in an increased R_h_.

**Figure 2.**
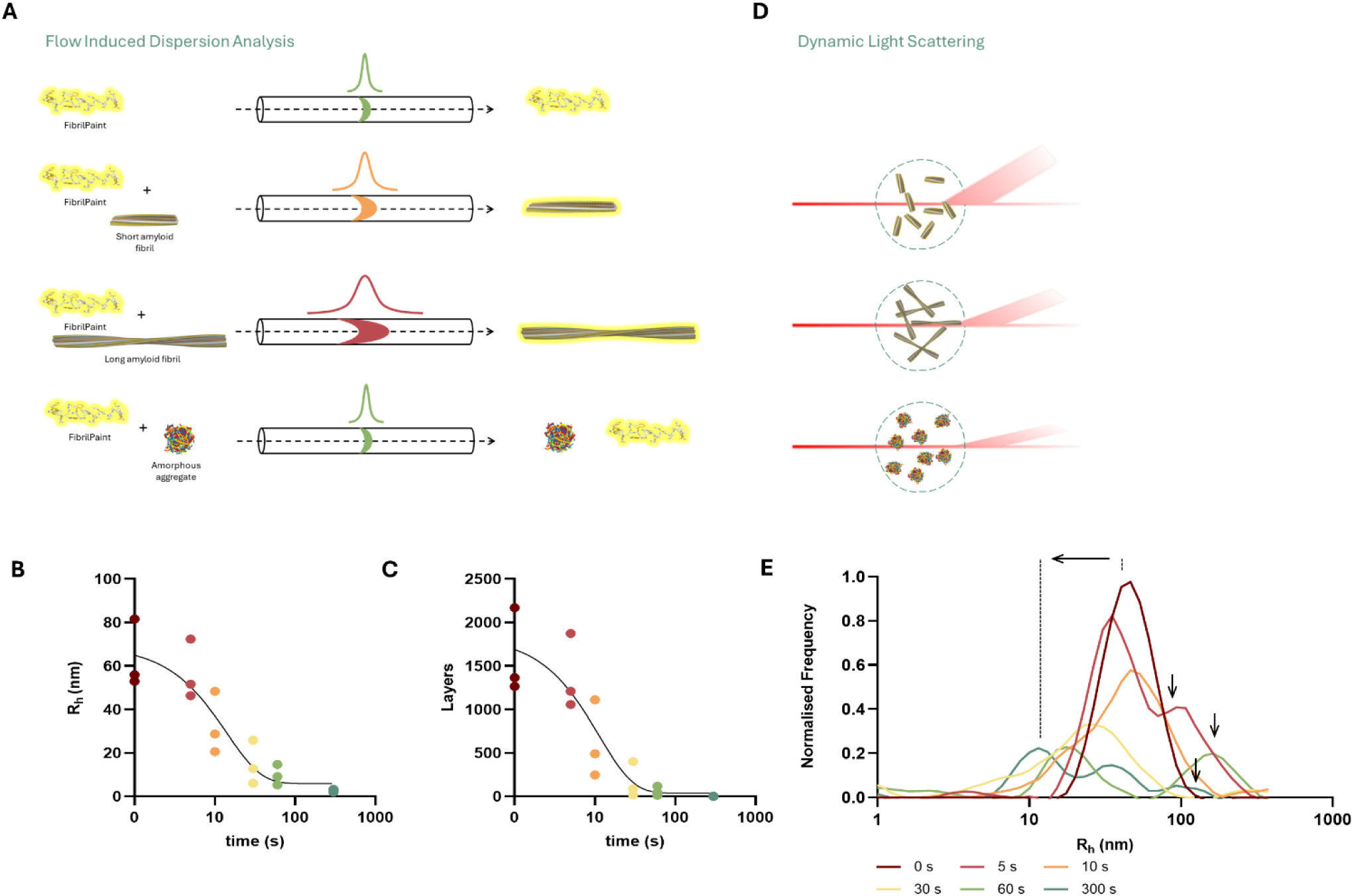
TauRD sonication reduces recombinant fibril size over time. Size analysis of TauRD fibril sonication at different time points. (A) Schematical representation of FP/FIDA assay. FP is specific of amyloid structures. FIDA calculates Rh based on diffusion in solution. FibrilPaint alone diffuses fast, resulting in a narrow peak and a small Rh, when bound to an amyloid species it will defuse slower, resulting in a broad peak and a big Rh. In presence of non-amyloid aggregates, FibrilPaint will stay unbound, diffuse fast and the Rh measured will be the same as FibrilPaint alone. (B) Rh of seeds measured using FibrilPaint in FIDA (half-time = 10.8 s). (C) Correlation to layers forming seeds. (D) Schematical representation of DLS. Based on scattered light related to size, DLS provides Rh of all particles regardless of structure and shape. (E) Rh of seeds measured in DLS.

Following this approach, we measured the size of seeds after direct sonication of pre-formed TauRD fibrils at different times. While FibrilPaint6 has an a R_h_ of 1.04 nm, the fluorescence signal increased to an average R_h_ of 64 nm in the presence of recombinant TauRD fibrils. Ultrasound fragmented these fibrils, strongly reducing the R_h_ value to 3 nm after 300 s of sonication (Fig. 2B, Sup. Fig. 1C). The half-breaking time of TauRD fibrils is 10.8 s. To better understand the effect of sonication on fibril size, we correlated R_h_ to layers forming the fibril and seeds. We have previously developed and application to predict fibril layers based on measured R_h_ (Sup. Fig. 1A, B)^20^, allowing us to have the relation between R_h_ and layers forming fibrils. Based on that, TauRD fibrils consist of 1601 layers, and after 300 s of sonication it is reduced to 2 layers (Fig 2 C, Sup. Fig. 1C).

Next, we further characterised the sonication products. Fibril Paint peptides exclusively interact with fibrils but no other aggregates ^20, 21^. Thus, we used Dynamic Light Scattering (DLS) to visualise also possible non-fibril sonication products. DLS provides the size of particles in solution by analysing the light scattering caused by the Brownian motion of those particles^24, 25^. As a result, it measures the R_h_ of all particles in solution, independent of their structure (Fig. 2D). We determined the R_h_ of TauRD sonicated samples. Upon increase of time, there is a shift to smaller R_h_ (Fig. 1E, Sup. Fig. 2). Interestingly, bigger species also appear (Fig. 1E, Sup. Fig. 2), which are only detectable in DLS. Since Fibril Paint is specific of amyloid structures, we can conclude that those are non-amyloid structures. Altogether, these results suggest that sonication breaks fibrils in smaller amyloid structures but also forms non-amyloid species.

### From 3150 to 80 layers in five minutes

Next, we investigated the effect of sonication in patient-derived fibrils. Those fibrils differ from the recombinant fibrils in structure^26, 27^. Additionally, patient fibrils have a fuzzy coat, lacking in TauRD fibrils, undergo post-translational modification and have different interfaces. We wanted to assess whether these different characteristics lead to different effect on the sonication. AD fibrils consist of PHF and Straight Filaments (SF), which are two identical C-shaped protofilaments with two different lateral contact leading to two different interfaces (Fig. 3A)^28^. After 300 s of sonication, the R_h_ of AD fibrils was reduced from 72 to 7 nm, with a half-breaking time of 2.8 s (Fig. 3B, Sup. Fig. 1D), which corresponds to a reduction from 1849 to 31 layers (Fig. 3C, Sup. Fig. 1D). CBD also contains two types of fibrils, type I and type II^29^ (Fig. 3A). Type I consist of a single protofilament and type II of a pair, which makes it the biggest fold. With a half-breaking time of 6 s, the size of CBD fibrils reduced from 145 nm to 7 nm after 300 s of sonication (Fig 3D, Sup. Fig. 1E). This R_h_ corresponds to a decrease from 4108 to 29 layers (Fig. 3E, Sup. Fig. 1E). The fibrils in FTD consist of a single protofilament^30^ (Fig. 3A), and the R_h_ after 300 s was reduced from 125 nm to 17 nm, and the half time was of 6.3 s (Fig 3F, Sup. Fig. 1F). The correlation to layers is from 3501 to 174 layers (Fig. 3G, Sup. Fig. 1E). Taken together, these results indicate that, independently of the differences in interface and structure, patient-derived fibrils are also susceptible to fragmentation.

**Figure 3.**
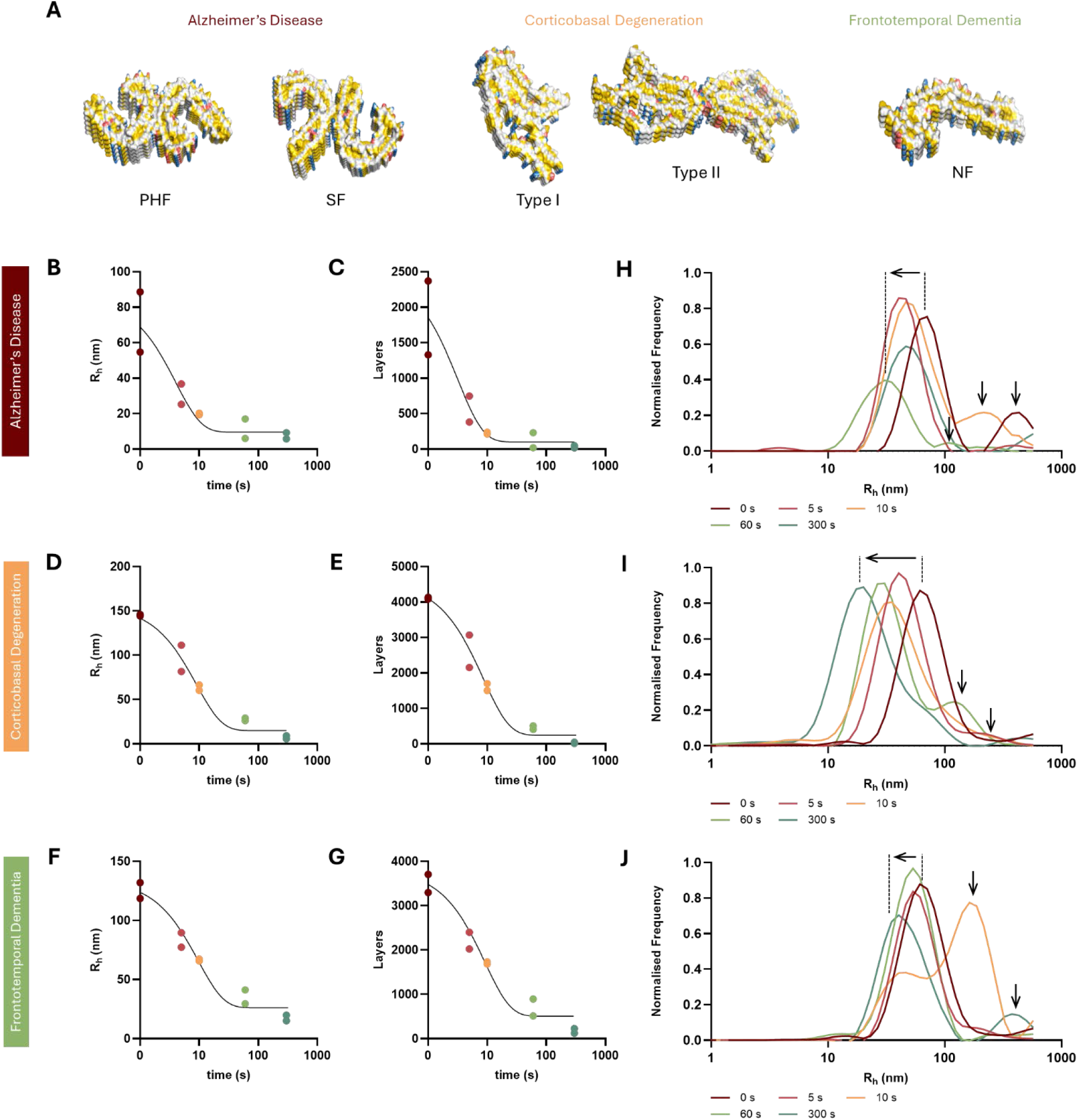
Patient derived Tau fibrils are susceptible to sonication. Size analysis of patient derived fibril sonication at different time points. (A) Structural representation of AD, CBD and FTD Tau fibrils (PHF, 5O3L; SF, 5O3T; Type I 6TJO; Type II 6VH7; NF 6GX5. Functional groups are coloured according to YRB script). (B) Rh of AD seeds measured using FP in FIDA (half-time = 2.8 s). (C) Correlation to layers forming AD seeds. (D) Rh of CBD seeds measured using FibrilPaint in FIDA (half-time = 6.03 s). (E) Correlation to layers forming CBD seeds. (F) Rh of FTD seeds measured using FibrilPaint in FIDA (half-time = 6.3 s). (G) Correlation to layers forming FTD seeds. (H) Rh of AD seeds measured in DLS. (I) Rh of CBD seeds measured in DLS. (J) Rh of FTD seeds measured in DLS.

After confirming the impact of sonication in amyloid size, we wanted to identify the presence of other type of aggregates. We measured the size of the sonicated patient fibrils using DLS. We observed similar results than the recombinant fibrils. While the R_h_ of AD, CBD and FTD decreases over sonication time, larger species also appear (Fig 3H-J, Sup. Fig 3-5). These results indicate that the sonication of AD, CBD and FTD fibrils leads to a combination of amyloid and non-amyloid species.

### Energy and fragmentation time of Tau fibrils

To understand the structural stability of the fibrils, we determined the required energy to fragmentate the different types of fibrils. Based on the parameters used to perform fibril sonication, we calculated the energy used among time (Fig. 4A). With the previously stablished the half-breaking time (Fig. 2B, C, Fig. 3B-G), we can determine the energy need to fragment in half the fibrils (Fig. 4B). AD fibrils need the lowest energy to be broken in half (840 J), followed by CBD and FTD, which need similar energy (1800 and 1890 J, respectively). Remarkably, TauRD fibrils, which have the smallest surface and only consist of one single protofilament, show the highest resistance to breaking, requiring the highest energy (3240 J) (Fig. 4C).

**Figure 4.**
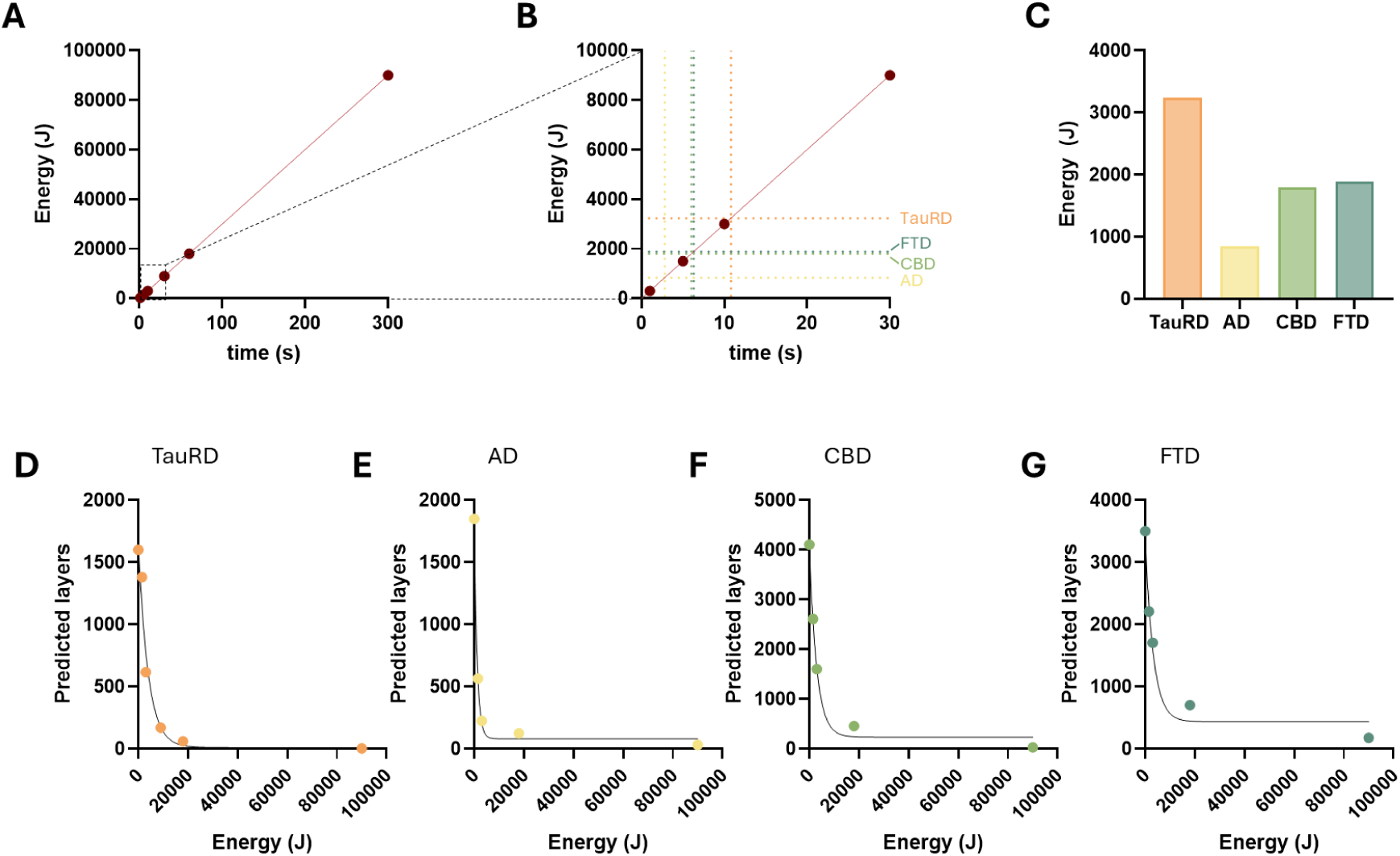
Quantification of energy in fibril fragmentation. (A) Energy used per time of sonication. (B) Zoom in of (A), dashed lines correspond to energy used in half-breaking time. (C) Energy needed in half-breaking time of Tau fibrils. Relation between energy and predicted layers during sonication of (D) TauRD, (E) AD, (F) CBD and (G) FTD.

We established the correlation between layers and energy (Fig. 4D-G), revealing an inverse relationship between these parameters, as expected. This relationship can be described by an exponential decay function. We can determine the precise value of energy required to achieve a specific length of sonicated fibrils.

## Discussion

Fibril fragmentation plays a significant role in amyloid aggregation by breaking fibrils in smaller seeds. Sonication is a commonly used method used to rapidly induce fibril fragmentation by applying high frequency sound waves that break the fibrils^11–16^. Here, we defined the impact of ultrasound on fibril fragmentation and the products resulting from it. We analysed the size of fibrils post-sonication using two different approaches. The first one, Fibril Paint/FIDA assay, measures the size of amyloid species, while the second approach, DLS, measures the size of all molecules. Both approaches give R_h_ in nm, allowing a quantitative description of sonication. We studied the effect of sonication using four different types of Tau fibrils, TauRD recombinant fibrils and patient-derived fibrils from AD, CBD and FTD. Notably, we observed that all types of fibrils are susceptible to sonication, despite these fibrils differ in structure and interface^26–30^.

Interestingly, we detected differences with the two approaches. The Fibril Paint/FIDA assay shows a time-dependent exponential decay in size, while in DLS, R_h_ reduces over time, but larger species also appear. This can be explained due to the differences in specificity of the two methods. The Fibril Paint/FIDA assay is specific for amyloid structures^20, 21^. FibrilPaint is a fluorescently labelled peptide that only binds to amyloid species, and only these will be measured by FIDA. In contrast, DLS measures Brownian motion of all molecules in solution, independent of their structure^24, 25^. This can explain that, as a result of fibril sonication, fibrils fractionate into smaller amyloid structures and non-amyloid structures, leading to a heterogeneous seed population.

Resistance to fibril fragmentation is associated to their stability, which can be influenced by multiple types of stress. Other methods to cause fibril fragmentation include mechanical agitation and thermal stress^32–35^. The effect of those methods, including sonication, on fibril fragmentation has been previously studied using various types of fibrils. Evidence suggests that fibril fragmentation exhibits increased toxicity ^32, 33^. Additionally, it has been reported that fibril fragmentation depends on polymorphism and preferential breaking positions ^16, 34, 35^, emphasising the complexity of this phenomenon. Altogether, understanding the mechanisms and conditions that dictate fibril fragmentation, can significantly contribute to unravelling the mechanism under amyloid-related diseases.

Importantly, the Fibril Paint/FIDA assay allows the quantification of fibril fragmentation. With it we can precisely determine the length, in layers forming fibrils, of the seeds resulting from sonication. Interestingly, our results show that sonication as a method to induce fibril fragmentation leads to a heterogeneous population of seeds, including amyloid and non-amyloid species. It will be interesting to further investigate their role in amyloid formation and associated toxicity. The hypothesised role of seeds in toxicity of neurodegenerative diseases^7–9^, positions them as a promising target for disease-modifying therapies. However, their complex nature represents a significant challenge. Understanding fibril fragmentation and stability may provide insight into the mechanism underlying amyloid formation, contributing to the development of therapeutic strategies targeting fibril stability and products resulting from fragmentation.

## Methods

### Expression and purification of TauRD

We produced N-terminally FLAG-tagged (DYKDDDDK) human TauRD (Q244-E372, with pro-aggregation mutation ΔK280) in E. Coli BL21 Rosetta 2 (Novagen), with an additional removable N-terminal His_6_-Smt-tag (MGHHHHHHGSDSEVNQEAKPEVKPEVKPETHINLKVSDGSSEIFFKIKKTTPLRRLMEAFAKRQGKEMDSLRFLYDGIRIQADQTPEDLDMEDNDIIEAHREQIGG). Expression was induced at OD_600_ 0.8 by addition of 0.15 mM IPTG and incubation at 18 °C overnight. Cells were harvested by centrifugation, resuspended in 25 mM HEPES-KOH pH=8.5, 50 mM KCl, flash frozen in liquid nitrogen, and kept at – 80 °C until further usage. Pellets were thawed at 37 °C, followed by the addition of ½ tablet/50 ml EDTA-free Protease Inhibitor and 5 mM β-mercaptoethanol. Cells were disrupted using an EmulsiFlex-C5 cell disruptor, and lysate was cleared by centrifugation. Supernatant was filtered using a 0.22 μm polypropylene filtered and purified with an ÄKTA purifier chromatography System. Sample was loaded into a POROS 20MC affinity purification column with 50 mM HEPES-KOH pH 8.5, 50 mM KCl, eluted with a linear gradient 0-100%, 5 CV of 0.5 M imidazole. Fractions of interest were collected, concentrated to 2.5 ml using a buffer concentration column (vivaspin, MWCO 10 KDa), and desalted using PD-10 desalting column to HEPES pH 8.5, ½ tablet/50 ml cOmplete protease inhibitor, 5 mM β-mercaptoethanol. The His_6_-Smt-tag was removed by treating the sample with Ulp1, 4 °C, shaking, overnight. The next day, sample was loaded into POROS 20HS column with HEPES pH 8.5, eluted with 0-100% linear gradient, 12 CV of 1M KCl. Fractions of interest were collected and loaded into a Superdex 26/60, 200 pg size exclusion column with 25 mM HEPES-KOH pH 7.5, 75 mM NaCl, 75 mM KCl. Fractions of interest were concentrated using a concentrator (vivaspin, MWCO 5 kDa) to desired concentration. Protein concentration was measured using a NanoDrop™ OneC UV/Vis spectrophotometer and purity was assessed by SDS-PAGE. Protein was aliquoted and stored at – 80 °C.

### Fibril extraction

Brain material of FTD and CBD were obtained from the Dutch Brain Bank, project number 1369. Brain material for AD was donated by prof. J. J. M. Hoozemans from the VU Medical Centra Amsterdam.

PHFs and SFs were extracted from grey matter of prefrontal cortex from one patient diagnosed with AD according to established protocols^28^. Tissue was homogenized using a Polytron(PT 2500E, Kinematica AG) on max speed in 20 % (w/v) A68 buffer, consisting of 20 mM TRIS-HCl pH 7.4, 10 mM EDTA, 1.6 M NaCl, 10% sucrose, 1 tablet/10 ml Pierce protease inhibitor, 1 tablet/10 ml phosphatase inhibitor. The homogenized sample was spun from 20 minutes, at 14000 rpm at 4 °C. Supernatant was collected, and the pellet was homogenized in 10% (w/v) A68 buffer. The homogenized was spun once more. The supernatants of both centrifugations were combined and supplied with 10% w/v Sarkosyl and incubated for 1 hour on a rocker at room temperature. The sample was ultracentrifuged for 1 hour at 100000xg and 4 °C. The supernatant was discarded, and the pellet was incubated overnight at 4 °C in 20 μl/0.2 g starting material of 50 mM TRIS-HCl pH 7.4. The next day, the pellet was diluted up to 1 ml of A68 buffer, and resuspended. To get rid of contamination, the sample was spun for 30 minutes at 14000 rpm, 4 °C. Supernatant was collected and spun once more for 1 hour at 100000 g, 4 °C. The pellet was resuspended in 30 μl 25 mM HEPES-KOH pH 7.4, 75 mM KCl, 75 mM NaCl, and stored at 4 °C up to a month. Presence of PHFs and SFs was assessed by Transmission Electron Microscope.

CBD fibrils were extracted from the grey matter of the superior parietal gyrus of one patient diagnosed with CBD (nhb 2018-007), following established protocols ^29^. Tissue was homogenized using a Polytron (PT 2500E, Kinematica AG) on max speed in 20 % w/v 10 mM TRIS-HCl pH 7.5, 1 mM EGTA, 0.8 M NaCl, 10% sucrose. The homogenate was supplied with 2% w/v of sarkosyl and incubated for 20 minutes at 37 °C. The sample was centrifuged for 10 minutes at 20000 g, and 25 °C. The supernatant was ultracentrifuged for 20 minutes at 100000 g and 25 °C. The pellet was resuspended in 750 μl/1g starting material of 10 mM TRIS-HCl pH 7.5, 1 mM EGTA, 0.8 M NaCl, 10% sucrose, and centrifuged at 9800 g for 20 minutes. The supernatant was ultracentrifuged for 1 hour at 100000 g. The pellet was resuspended in 25 μl/g starting material of 20 mM TRIS-HCl pH 7.4, 100 mM NaCl, and stored at 4 °C up to a month. Presence of CBD fibrils was assessed by Transmission Electron Microscope.

Narrow filaments were extracted from the grey matter of the middle frontal gyrus from one patient diagnosed with FTD (nhb 2017-019), as described in established protocols^30^. Fibrils were extracted following the protocol for AD fibrils. After the first ultracentrifugation step, the pellet was resuspended in 250 μl/1g of starting material of 50 mM Tris pH 7.5, 150 mM NaCl, 0.02% amphipol A8-35. The sample was centrifuged for 30 minutes at 3000 g and 4 °C. Pellet was discarded, and the supernatant was ultracentrifuged for 1 hour at 100000 g and 4 °C. The pellet was resuspended in 30 μl of 50 mM TRIS-HCl pH 7.4, 150 mM NaCl, and stored at 4 °C up to a month. Presence of narrow filaments was assessed by Transmission Electron Microscope.

### Peptide synthesis and purification

The peptides were synthesized using a Liberty Blue MicrowaveAssisted Peptide Synthesizer (CEM) with standard Fmoc chemistry and DIEA/HBTU as coupling reagents. The peptide concentrations were measured by UV spectroscopy. The peptides were labelled with 5(6)-carboxyfluorescein at their N’ termini^36^. The peptides were cleaved from the resin with a mixture of 95 % (v/v) trifluoroacetic acid (TFA), 2.5% (v/v) triisopropylsilane (TIS), 2.5 % (v/v) triple distilled water (TDW) agitating vigorously for 3 hours at room temperature. The volume was decreased by N_2_ flux, and the peptides precipitated by addition of 4 volumes of diethylether at −20 °C. Thepeptides were sedimented at −20 °C for 30 minutes, then centrifuged and the diethylether discarded. The peptides were washed three times with diethylether and dried by gentle N_2_ flux. The solid was dissolved in 1:2 volume ratio of acetonitrile (ACN):TDW, frozen in liquid Nitrogen and lyophilized. The peptides were purified on a WATERS HPLC using a reverse-phase C18 preparative column with a gradient ACN/TDW. The identity and purity of the peptides was verified by MALDI mass spectrometry and Merck Hitachi analytical HPLC using a reverse-phase C8 analytical column.

### TauRD fibril formation

Aggregation of 20 μM TauRD in 25 mM HEPES-KOH pH 7.4, 75 mM KCl, 75 mM NaCl, ½ tablet/50 ml Protease Inhibitor, was induced by the addition 5 μM of heparin low molecular weight and incubated at 37 °C and 600 rpm in an EppendorfTM Thermomixer® during 20 hours. For ThT aggregation assay 45 μM of ThT were added to the reaction, and fluorescent spectra was recorder every 5 minutes, at 37 °C and 600 rpm during 24 hours in CLARIOstar® Plus.

### Seeds preparation

Tau fibrils were diluted to a final concentration of 2.5 μM in 25 mM HEPES-KOH pH 7.5, 75 mM KCl, 75 mM NaCl, 0.5% pluronic (for TauRD and AD), 20 mM TRIS-HCl pH 7.4, 100 mM NaCl (for CBD) or 50 mM TRIS-HCl pH 7.4, 150 mM NaCl (for FTD), in a total volume of 50 μl. Samples were sonicated using a Sonics VibraCell at 40% amplitude, 1 s on and 1 s off during the desired time.

### Size measurement with FIDA

2 μM of fibrils or seeds size was measured in presence of 200 nM of FibrilPaint was measured in 25 mM HEPES-KOH pH 7.5, 75 mM KCl, 75 mM NaCl, 0.5% pluronic (for TauRD and AD), 20 mM TRIS-HCl pH 7.4, 100 mM NaCl, 0.5% pluronic (for CBD) or 50 mM TRIS-HCl pH 7.4, 150 mM NaCl, 0.5% pluronic (for FTD). Size measurements were performed in a FIDA1 instrument with a 480 nm excitation source, using a capillary dissociation approach (Table 1, 2).

**Table 1.**
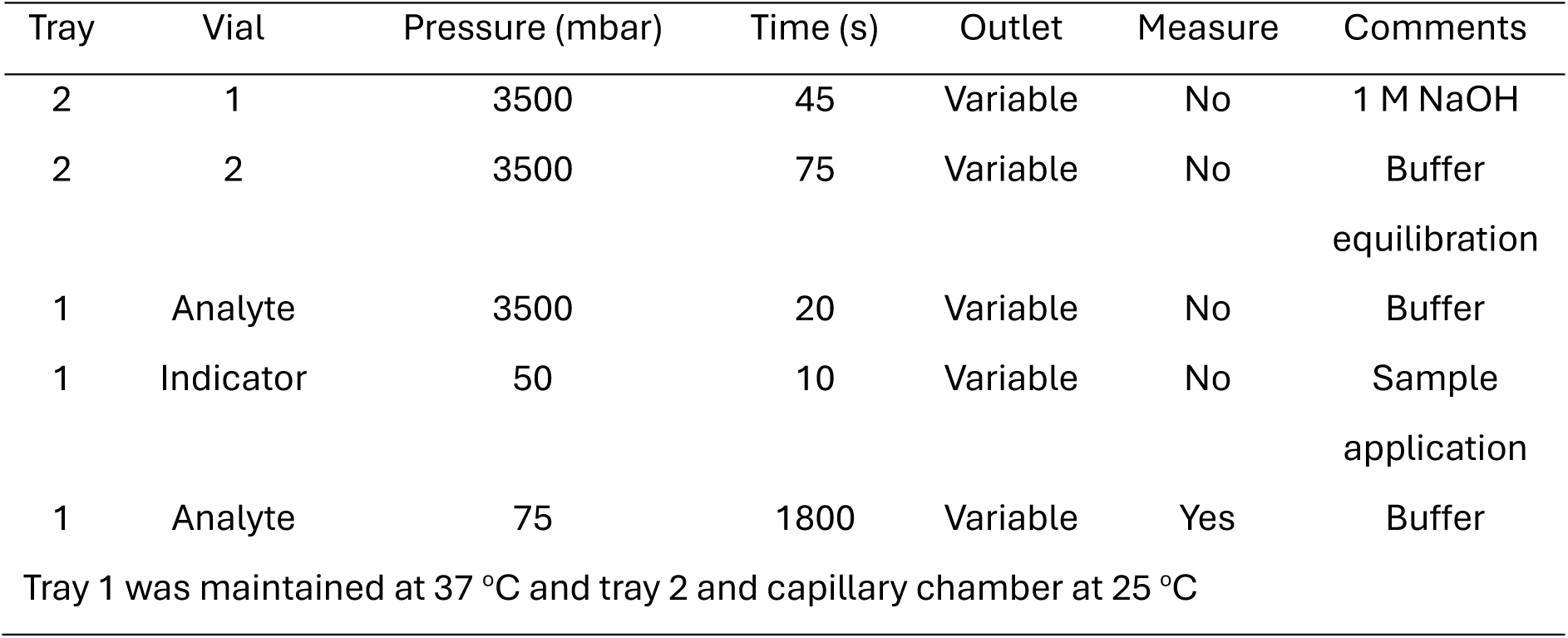
Experimental parameters for analysis of proteins up to R_h_ of 75 nm.

**Table 2.** Experimental parameters for analysis of proteins up to R_h_ of 150 nm.

| Tray | Vial | Pressure (mbar) | Time (s) | Outlet | Measure | Comments |
| --- | --- | --- | --- | --- | --- | --- |
| 2 | 1 | 3500 | 45 | Variable | No | 1 M NaOH |
| 2 | 2 | 3500 | 75 | Variable | No | Buffer<br>equilibration |
| 1 | Analyte | 3500 | 20 | Variable | No | Buffer |
| 1 | Indicator | 50 | 10 | Variable | No | Sample<br>application |
| 1 | Analyte | 30 | 5000 | Variable | Yes | Buffer |

Tray 1 was maintained at 37 °C and tray 2 and capillary chamber at 25 °C

### Size measurement with DLS

Fibrils or seeds were diluted to a final concentration of 2 μM in 25 mM HEPES-KOH pH 7.5, 75 mM KCl, 75 mM NaCl (for TauRD and AD), 20 mM TRIS-HCl pH 7.4, 100 mM NaCl (for CBD) or 50 mM TRIS-HCl pH 7.4, 150 mM NaCl (for FTD) and loaded into High Sensitivity glass capillaries. Measurements were performed at 37 °C, 100% excitation power and 100% DLS laser power.

### R_h_ and layer prediction model

We generated PHF (53OL) consisting of different lengths by assembling a defined number of layers in a stack configuration. We used those structures to predict their R_h_ using a FIDAbio R_h_ prediction tool. Due to minimal size differences among other fibril types, the PHF structure was used in all cases to predict size.

## Supplementary Figures

**Supplementary Figure 1.**
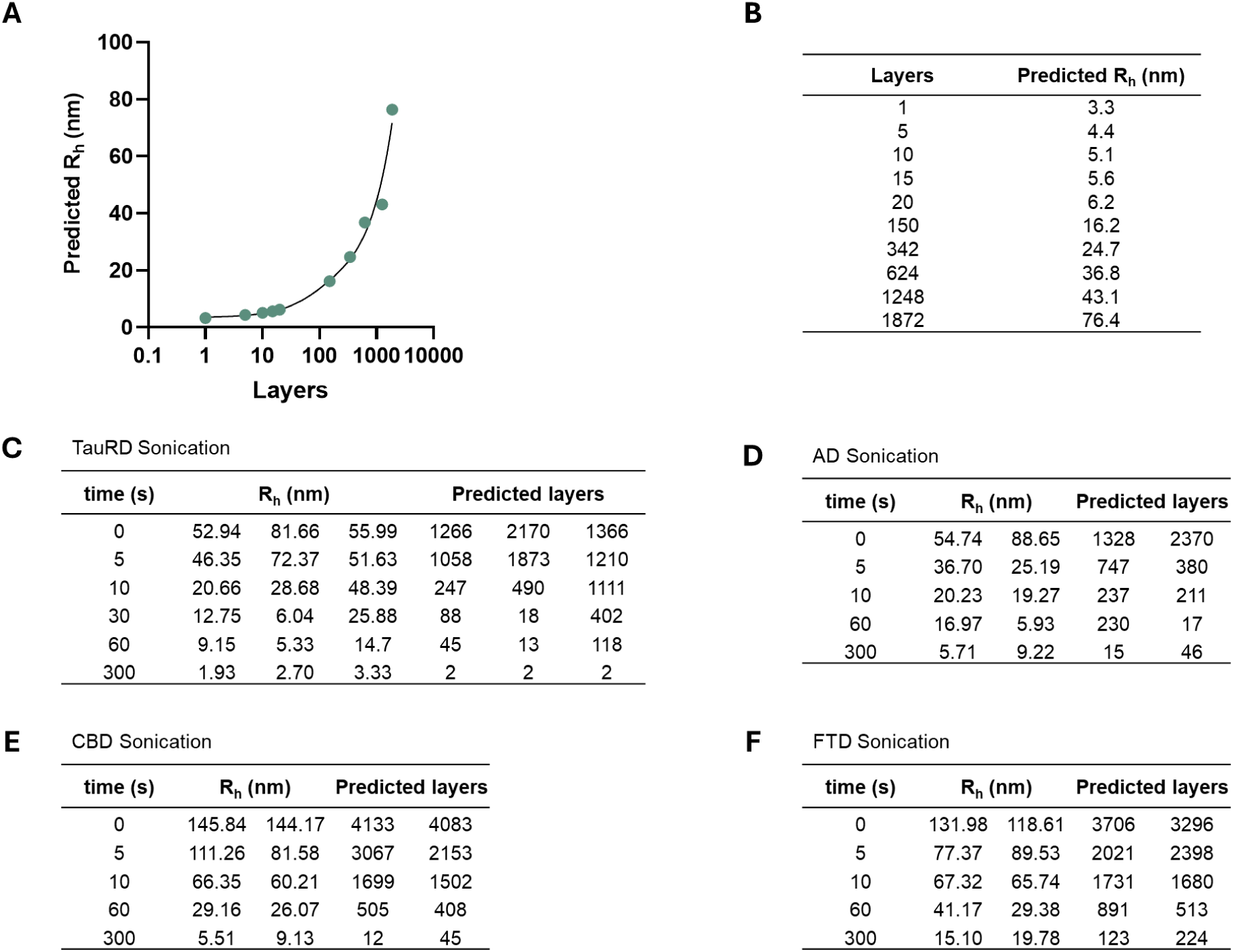
Correlation Rh to PHF layers. (A) Rh prediction of AD PHF of different layers using FIDAbio prediction tool. (B) Experimental values used in (A). Measured Rh and predicted layers of (C) TauRD fibrils (D) AD fibrils (E) CBD fibrils and (F) FTD fibrils sonication at different times.

**Supplementary Figure 2.**
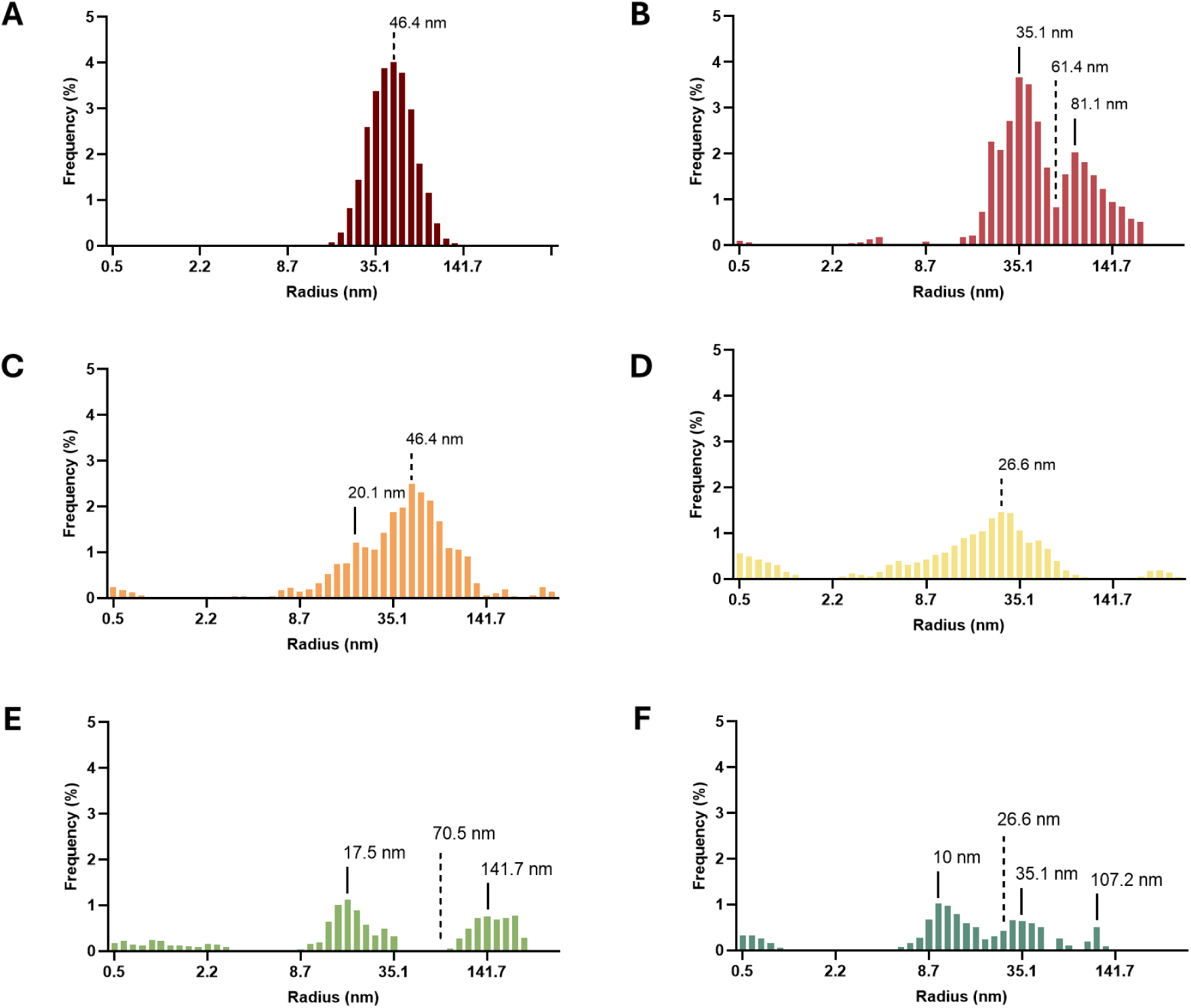
DLS analysis of TauRD fibrils sonication over time. Analysis of radius of TauRD fibrils after (A) 0 s, (B) 5 s, (C) 10 s, (D) 30 s, (E) 60 s and (F) 300 s of sonication. Dashed line indicates cumulant radius.

**Supplementary Figure 3.**
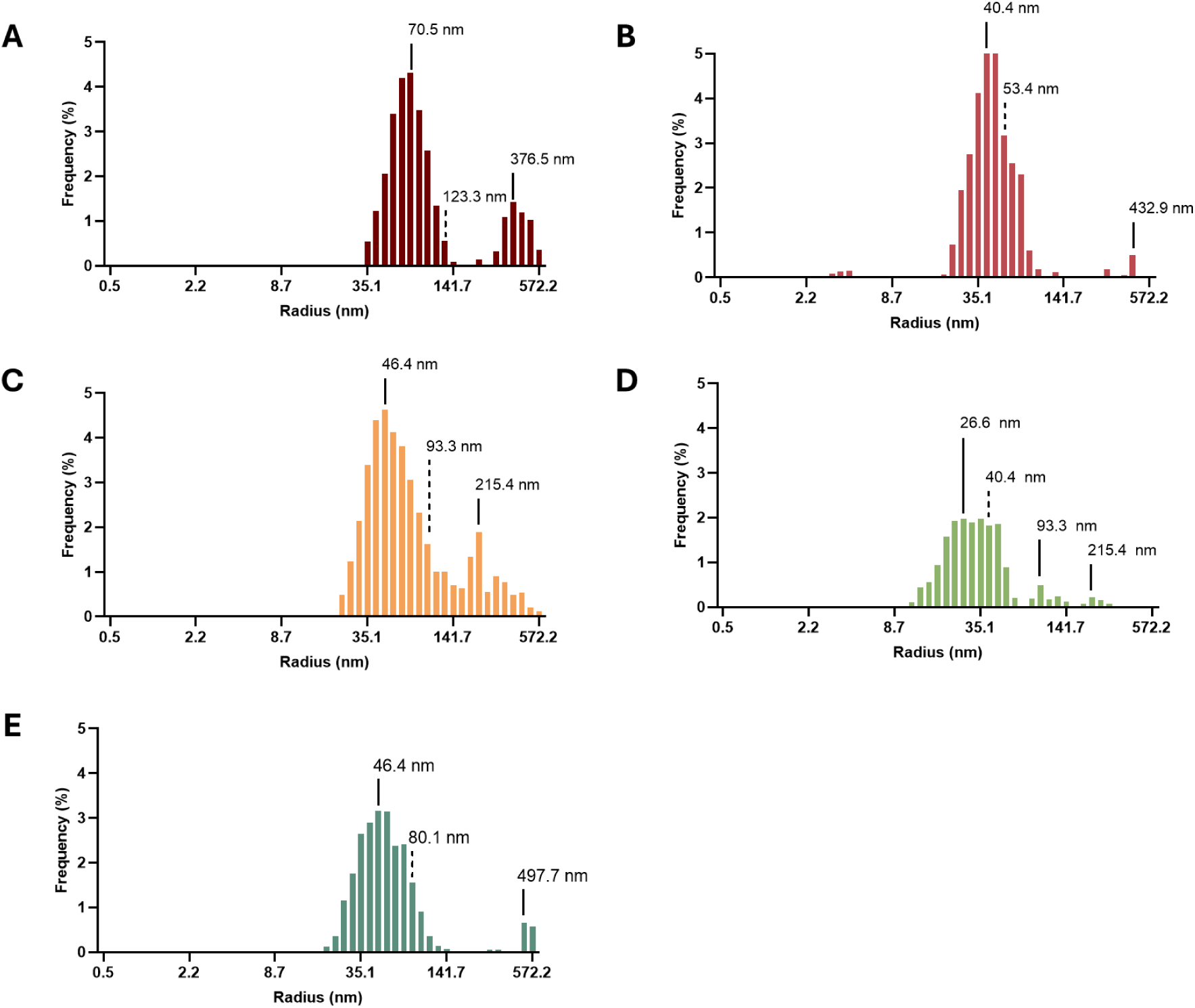
DLS analysis of AD fibrils sonication over time. Analysis of radius of AD fibrils after (A) 0 s, (B) 5 s, (C) 10 s, (D) 60 s and (E) 300 s of sonication. Dashed line indicates cumulant radius.

**Supplementary Figure 4.**
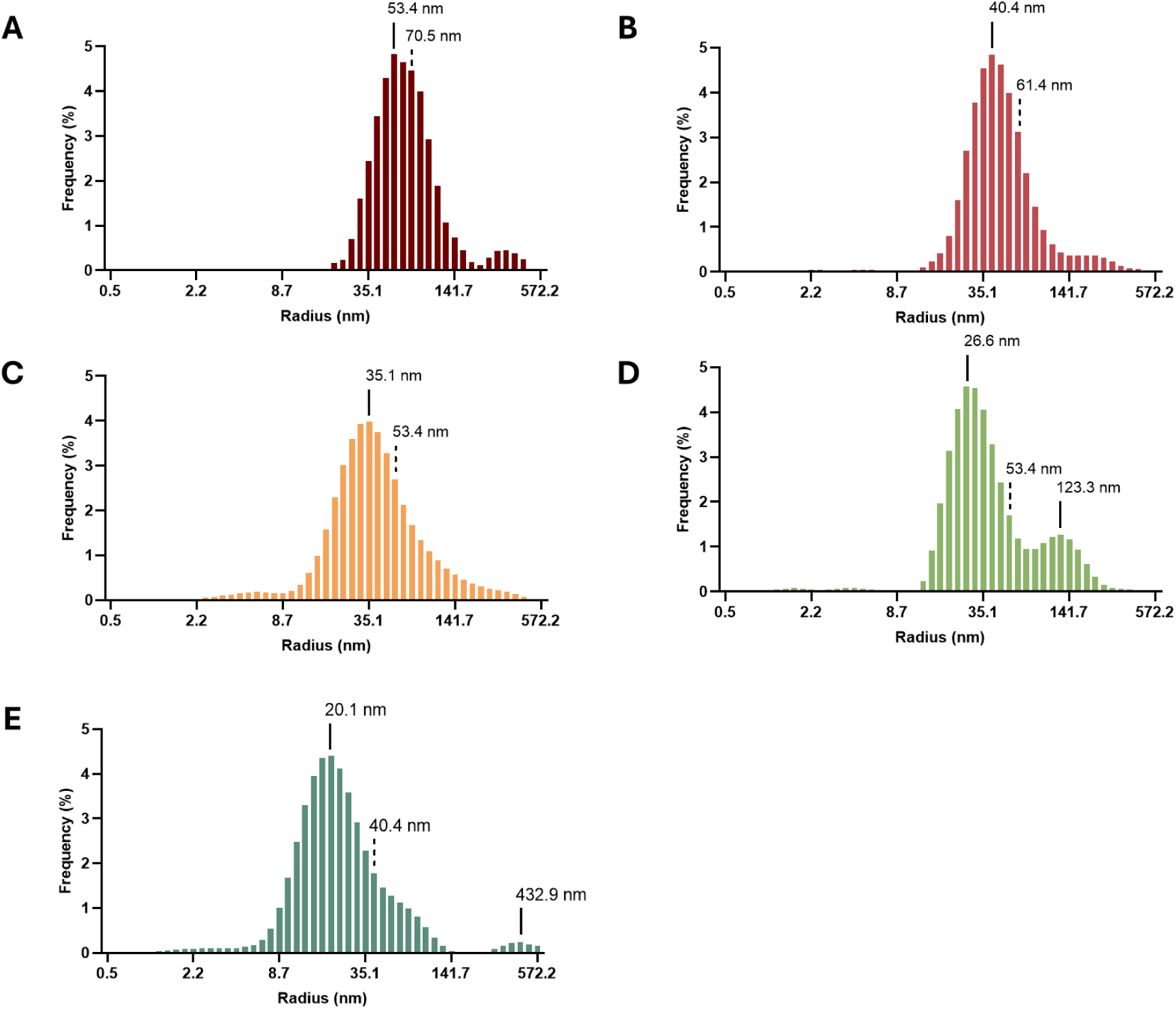
DLS analysis of CBD fibrils sonication over time. Analysis of radius of CBD fibrils after (A) 0 s, (B) 5 s, (C) 10 s, (D) 60 s and (E) 300 s of sonication. Dashed line indicates cumulant radius.

**Supplementary Figure 6.**
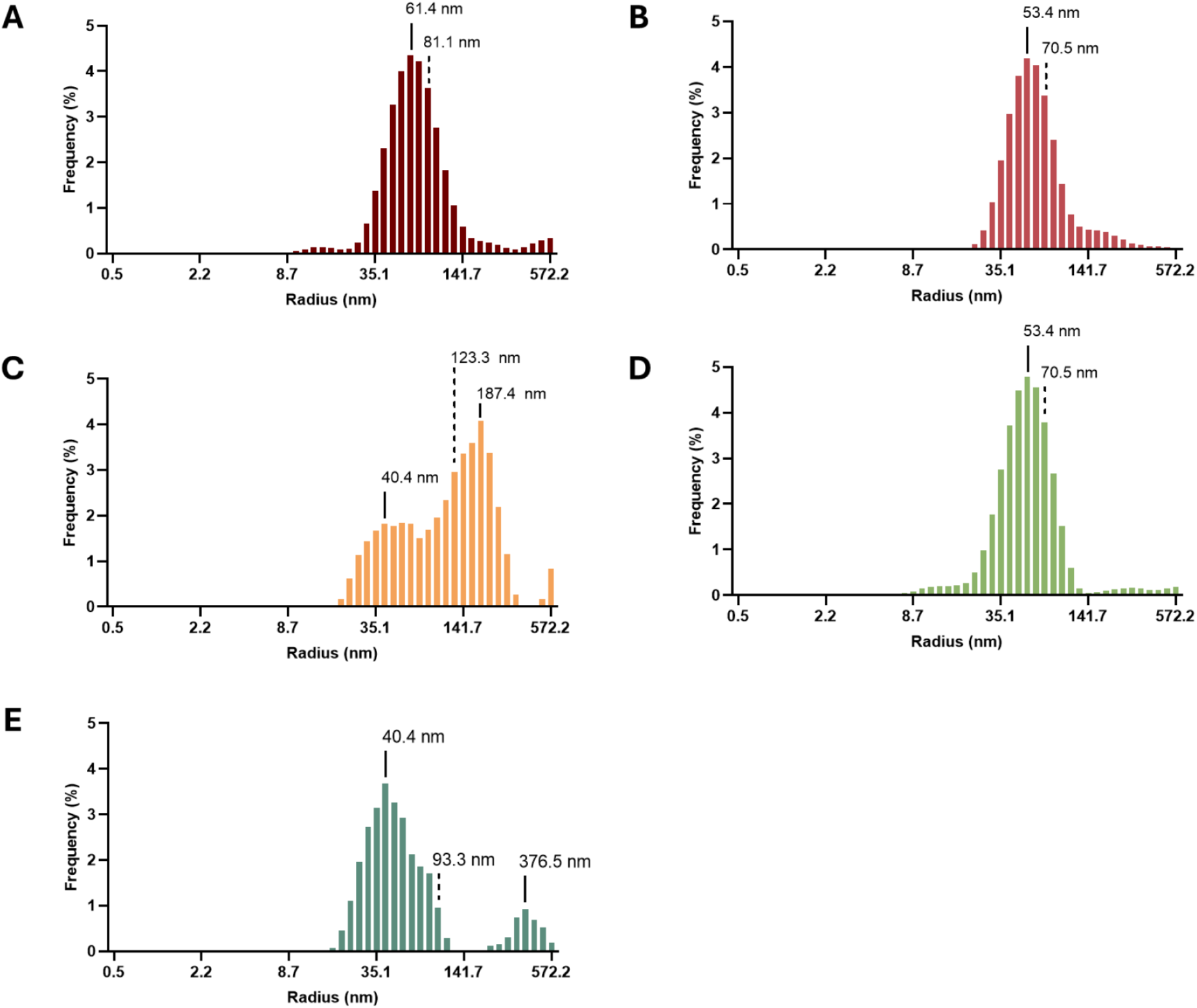
DLS analysis of CBD fibrils sonication over time. Analysis of radius of CBD fibrils after (A) 0 s, (B) 5 s, (C) 10 s, (D) 60 s and (E) 300 s of sonication. Dashed line indicates cumulant radius.

## Acknowledgements

SGDR was supported by grants of the Campaign Team Huntington and Alzheimer Nederland (No. WE.03-2019-03). SGDR acknowledges funding by the Zwaartekracht Grant FLOW, funded by the Netherlands Ministry for Education, Culture and Science. AF thanks the Minerva Center for Bio-Hybrid complex systems and the Saerree K. and Louis P. Fiedler Chair in Chemistry. Measurements on the CLARIOstar Plus®, FIDA1 and Prometheus Panta were done at the Protein Research Centre of Utrecht University.

## Author contribution

Conception JAP, SGDR

Design of the work JAP, SGDR

Acquisition JAP, MT, SS

Analysis JAP, MT, SGDR

Interpretation of data JAP, MT, SGDR

Writing original draft JAP

Revision JAP, AF, SGDR

